# Broad generalisation of the ventriloquism aftereffect across sound frequencies

**DOI:** 10.1101/2021.12.22.473801

**Authors:** Rachel Ege, Nina C. Haukes, A. John van Opstal, Marc M. van Wanrooij

**Author notes:** Shared first authorship (Rachel Ege, Nina C. Haukes).

## Abstract

Humans localise sounds in the horizontal plane by processing level and timing differences between the ears. This neurocomputational process is continuously and adaptively calibrated using visual input, as seen in the ventriloquism after effect: a shift in sound perception toward a previously seen light. It is unknown from where in the brain this aftereffect originates; adaptation could occur at an early level in the auditory system where neurons are narrowly tuned to frequency, at a later level in the auditory system where localisation cues are extracted, or outside the auditory system at a higher-level spatial map. To investigate this, we examined how the ventriloquism aftereffect generalises across sound frequencies. Participants localised seven narrowband sounds (0.5-8 kHz), targeting different localisation cues. We found that sound localisation accuracy in darkness varied slightly with frequency. When sounds were paired with a visual stimulus that was offset by 10 deg, participants exhibited a pronounced bias toward the light of about ∼63%, corresponding to the well-known ventriloquism effect. The bias was stronger for narrowband compared to broadband sounds. After exposure to a block of these audiovisual stimuli, a ventriloquism aftereffect in the form of a spatial bias of ∼12% was observed across all tested frequencies, largely independent of the frequency of the exposure sound. Together with earlier reports of both frequency-specific and frequency-general recalibration, our results indicate that under conditions of a fixed and consistent audiovisual spatial offset, the ventriloquism aftereffect generalises across sound frequencies, consistent with adaptation at a frequency-independent multisensory spatial stage.

## Introduction

Horizontal sound localisation relies on processing of binaural localisation cues^1,2^: interaural time differences (ITDs), which result from differences in sound arrival time between the ears and are prominent at low frequencies (≲1.5 kHz), and interaural level differences (ILDs), which arise from the acoustic shadow of the head and are prominent at higher frequencies (≳2-3 kHz). To maintain accurate auditory spatial perception, the mapping between these cues and external space must be continuously calibrated. Vision can serve as a calibration signal, which is illustrated by the ventriloquism aftereffect. After repeated exposure to a sound and a spatially displaced light, the perceived sound location remains biased toward the previous visual input, even in the absence of the light^3-6^. The ventriloquism aftereffect has been widely documented^7-18^, yet it remains unclear how this effect generalises. Specifically, it is unknown whether the aftereffect generalises across different frequencies, across binaural localisation cues (ITDs and ILDs), or across a higher-order spatial representation. Examining how the ventriloquism aftereffect behaves across spectral conditions can help distinguish between these possibilities. In this study, we assess how the spectral content of sounds influences the strength and generalisation of the aftereffect.

Previous research^4-6,9,13^ suggests that the ventriloquism aftereffect could originate from different levels of auditory processing, leading to three competing hypotheses (schematically illustrated in Fig. 1): (i) the frequency hypothesis, (ii) the cue hypothesis, and (iii) the space map hypothesis. The three mechanisms describe different levels in auditory processing. First, the frequency hypothesis (Fig. 1a) states that the audiovisual interactions between the sound and the light take place at a level where the auditory input is tonotopically represented. If the audiovisual interactions take place at this level, the ventriloquism aftereffect is expected to only occur for sound frequencies that lie within the critical spectral band around the exposure frequency^19,20^ (Fig. 1a, right). This would mean there will be no generalisation of the aftereffect to other sound frequencies (support for this hypothesis^4,5,13^).

**Figure 1.**
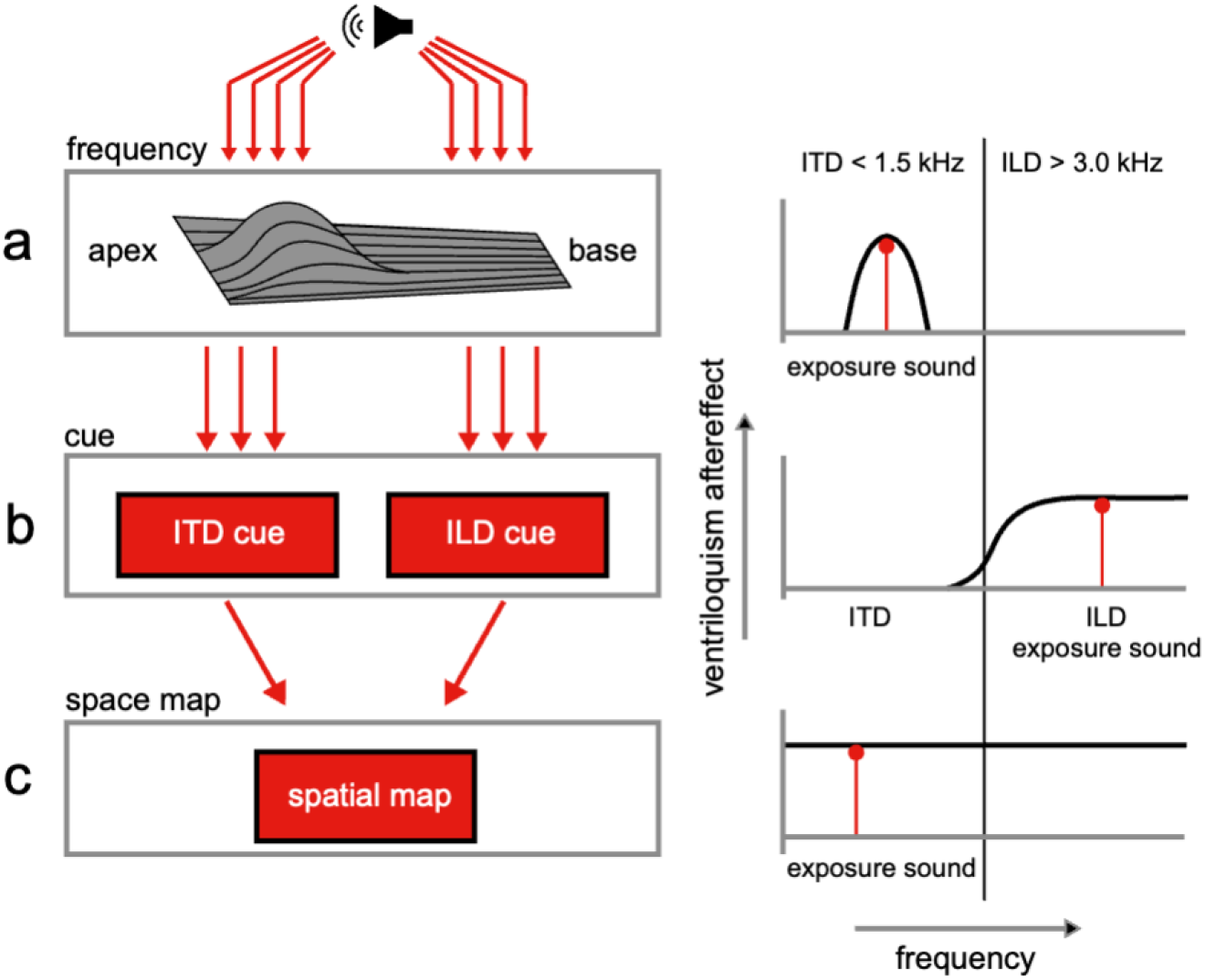
Ventriloquism aftereffect models. **a)** The influence of the light on sound localisation takes place at a tonotopic stage (left column). The ventriloquism aftereffect is then only found for frequencies within a critical band (right column, black curve) around the exposure frequency (right column, red vertical line with dot on the top). **b)** The light influences sound localisation when the ITD or ILD cues have been extracted (left column). The ventriloquism aftereffect will be restricted to frequencies belonging to that cue (right column, the black curve for ILD cues, given the frequency of the exposure sound, in red). The thin vertical line represents the border between the ITD and ILD domains (around 2 kHz). **c)** Generalisation across the entire frequency domain occurs if the light influences sound localisation at the level of a spatial map (left column). The aftereffect will generalise across all frequencies (left column, black line). The schematic depicts limiting cases; in practice, different mechanisms may dominate depending on experimental conditions.

Second, the cue hypothesis (Fig. 1b) states that the audiovisual interactions take place at a stage in auditory processing where the binaural spatial cues have been extracted. Broadband sounds, containing a wide range of frequencies, can be localised using both cues, whereas sounds with only frequencies between 1.5 and 3 kHz provide weak or ambiguous localisation cues. By using narrowband sounds with different frequencies, we can target different localisation cues (low frequency ITDs, high frequency ILDs). If the cue hypothesis holds, the ventriloquism aftereffect will generalise among all sound frequencies that share the same localisation cue as the exposure sound, but not to sound frequencies that rely on the other localisation cue (Fig. 1b, right; to our knowledge, the only study that directly tested this prediction found no evidence in support of it ^9^).

Third, the space map hypothesis states that audiovisual interactions occur at a multisensory or motor stage where auditory cues have been integrated with non-auditory cues, for example at an eye-centred motor map^2,12,21^ (Fig. 1c). If this holds, the ventriloquism aftereffect will generalise across all sounds (Fig. 1c, right; support for this hypothesis^6,9^).

Alternatively, the degree of generalisation may not depend on the stage of auditory processing, but rather on how auditory neurons across the auditory system are tuned to frequency as a function of sound intensity^22,23^. At high sound intensities, auditory neurons tend to respond to a broad range of frequencies, whereas at lower intensities, their frequency tuning becomes narrower. If this intensity hypothesis holds, the ventriloquism aftereffect should generalise fully across frequencies only for high sound levels (exceeding 66 dB according to simulations^23^). At lower intensities, the aftereffect is expected to be limited or absent due to more selective frequency tuning. This account could reconcile discrepancies between existing studies supporting frequency-specific (tonotopy) versus frequency-general (space map) hypotheses, due to different test configurations.

In this study, we used an absolute pointing task based on head movements to quantify how participants localised sounds across three experimental phases, conducted over three separate sessions. Each session featured a different exposure sound, either a low-frequency, high-frequency, or broadband sound, paired with a spatially displaced visual stimulus. In the pre-exposure phase, we assessed baseline localisation accuracy for eight sounds (one broadband and seven narrowband) presented in isolation, without any visual stimulus. In the exposure phase, we measured the immediate ventriloquism effect - a shift in perceived sound location caused by a visual stimulus offset by 10° - for the session-specific exposure sound. Finally, in the post-exposure phase, we assessed the ventriloquism aftereffect by measuring localisation responses to all eight sounds again, now without any visual input. This design allowed us to test whether exposure to a single sound leads to a lasting recalibration of sound localisation. All sounds were presented at a relatively low intensity (55 dB). This is a sound level at which Magosso et al. ^23^ predict minimal or no generalisation of the aftereffect, based on prior findings^4,5^ of limited generalisation at low intensities (45 and 60 dB), in contrast to studies at higher intensities (66 and 70 dB) that report generalisation^6,9^.

By measuring sound-localisation biases across a wide range of test frequencies following exposure to different sounds, the present design allows us to evaluate which predicted patterns of generalisation best describe the data. We find that the ventriloquism aftereffect generalises broadly across the tested frequency range: the post-exposure bias is neither confined to the exposure frequency nor restricted to sounds relying on the same binaural cue as the exposure stimulus. Instead, the aftereffect is expressed across all tested frequencies, consistent with recalibration at a frequency-independent, multisensory spatial stage under the present experimental conditions.

## Materials and Methods

### Participants

Twelve human participants provided their informed consent and performed the experiments. All participants had normal or corrected-to-normal vision and normal hearing (verified by means of a sound localisation experiment including low-pass, high-pass and Gaussian white noise sounds in the frontal hemifield, and a standard audiogram). Three participants helped in the execution of the study; the other participants were kept naive about the purpose of the study. The experimental protocol was approved by local institutional ethical committee of the Faculty of Social Sciences at the Radboud University (ECSW 2016-2208-41), and all applied procedures were in line with the relevant guidelines and regulations.

### Apparatus

During the experiment, the participant sat in a chair in the centre of a completely dark, sound-attenuated room. The room had an ambient background noise level below 30 dBA. The chair was positioned at the centre of a spherical frame (radius 1.5 m), on which 125 small broad-range loudspeakers (SC5.9; Visaton GmbH, Haan, Germany) were mounted. Green LEDs (wavelength 565 nm; Kingsbright Electronic Co., LTD., Taiwan; luminance 1.4 cd/m^2^) were mounted at the centre of each speaker. For a more thorough description of the room, see ^24^. Head movements were recorded with the magnetic search-coil technique^25^. To this end, the participant wore a lightweight spectacle frame with a small coil attached to their nose bridge. The horizontal and vertical head-coil signals were amplified and demodulated (EM7; Remmel Labs, Katy, TX, USA), low-pass-filtered at 150 Hz (custom built, fourth-order Butterworth), digitized (RA16GA and RA16; Tucker-Davis Technology, Alachua, FL, USA) and stored on hard disk at 6 kHz/channel. A custom-written MATLAB program, running on a PC (HP EliteDesk, California, United States) controlled data recording and storage, stimulus generation, and online visualization of the recorded data.

### Stimuli

Sounds were pre-generated as white noise. Seven narrowband sounds were created by band-pass filtering to have a bandwidth of 2/3 octaves around centre frequencies {500, 793, 1260, 2000, 3175, 5040, 8000} Hz. A broadband sound was generated by band-pass filtering between 0.5 and 20 kHz. All sounds had a duration of 75 ms, with 5-ms sine-squared onset, and cosine-squared offset ramps, and were presented at an A-weighted level of L_A_ = 55 dB.

### Calibration

To determine the off-line calibration that maps the raw coil signals (in mV) onto known target locations (in deg), participants pointed their head towards 24 LED locations in the frontal hemifield (separated by approximately 30 deg in both azimuth and elevation). The pointing was done via a red LED that was attached to the spectacle frame, at a fixed distance of 40 cm in front of the eyes. This setup ensured the position of the eyes was aligned with the head movements. A three-layer neural network (implemented in MATLAB) was trained to map the raw initial and final head position signals onto the (known) LED azimuth and elevation angles, with an overall accuracy of 1.0 deg or better. The weights of the neural network were subsequently used to map all head-movement voltages of the localisation experiments to degrees.

### Paradigm

Participants were instructed to localise sounds. Each participant performed three sessions, each on a separate day. A brief overview of each session is shown in Table 1. Each session consisted of three phases: pre-exposure, exposure, and post-exposure. During the pre- and post-exposure phases we presented eight sounds. During the exposure phase, one sound was presented in combination with an offset visual stimulus. This sound was either a 793 Hz, a 5040 Hz, or a broadband sound, using a different sound for each session. These stimuli were chosen to target specific localisation cues; a low-frequency narrowband sound to target ITDs, a high-frequency narrowband sound to target ILDs, and a broadband sound to target both cues. In the pre-exposure phase, we measured sound localisation performance, in the exposure phase we determined the ventriloquism effect, and in the post-exposure phase we obtained the ventriloquism aftereffect.

**Table 1.** Overview of stimulus conditions per session. Rows correspond to the three experimental phases within a session (pre-exposure, exposure, and post-exposure). Columns list the sounds presented (NB: narrowband, with 793 and 5040 Hz specified; BB: broadband), the horizontal target locations, the presence and spatial offset of a visual stimulus, the number of repetitions per sound-location combination (# presentations), and the resulting total number of trials per phase (# trials).

| Phase | Sounds Presented | Target Locations | Visual Stimulus | # Presentations | # Trials |
| --- | --- | --- | --- | --- | --- |
| Pre-exposure | 7 NB+1BB | -35° to +35° (10° steps;<br>8 locations) | None | 4 | 256 |
| Exposure | 1 sound (793, or 5040, or BB) | -35° to +35° | +10° offset | 56 | 448 |
| Post-exposure | 7 NB+1BB | -35° to +35° | None | 4 | 256 |

The design of the pre- and post-exposure phases were the same. Target sounds were presented at eight locations in the horizontal plane (-35 to +35 deg, in 10 deg steps). Each stimulus-location combination was presented four times in pseudo-random order (8 sounds x 8 locations x 4 presentations = 256 trials). During the exposure phase, one of three sounds was presented at the same eight locations (-35 to +35 deg, in steps of 10 deg), while simultaneously a LED was presented ten degrees to the right (- 25 to +45 deg, in steps of 10 deg). Each stimulus-location combination was presented 56 times, resulting in 448 trials (56 presentations x 8 locations). In each trial, participants first aligned the head-fixed laser with a central fixation light (Fig. 2). A fixation light was presented at the centre speaker (0 deg azimuth and 0 deg elevation) at the start of each trial, to ensure participants had a similar starting position over trials. The fixation light was extinguished with a random timing between 300-800 ms. After the fixation light there was a gap of 200 ms before the target stimulus was presented. Participants were asked to align the head-fixed laser pointer as quickly and as accurately as possible with the sound source’s origin (neglecting the light). One session took approximately 60 minutes, which includes the calibration and short breaks in between the blocks. The order of the sessions was randomised among participants.

**Figure 2.**
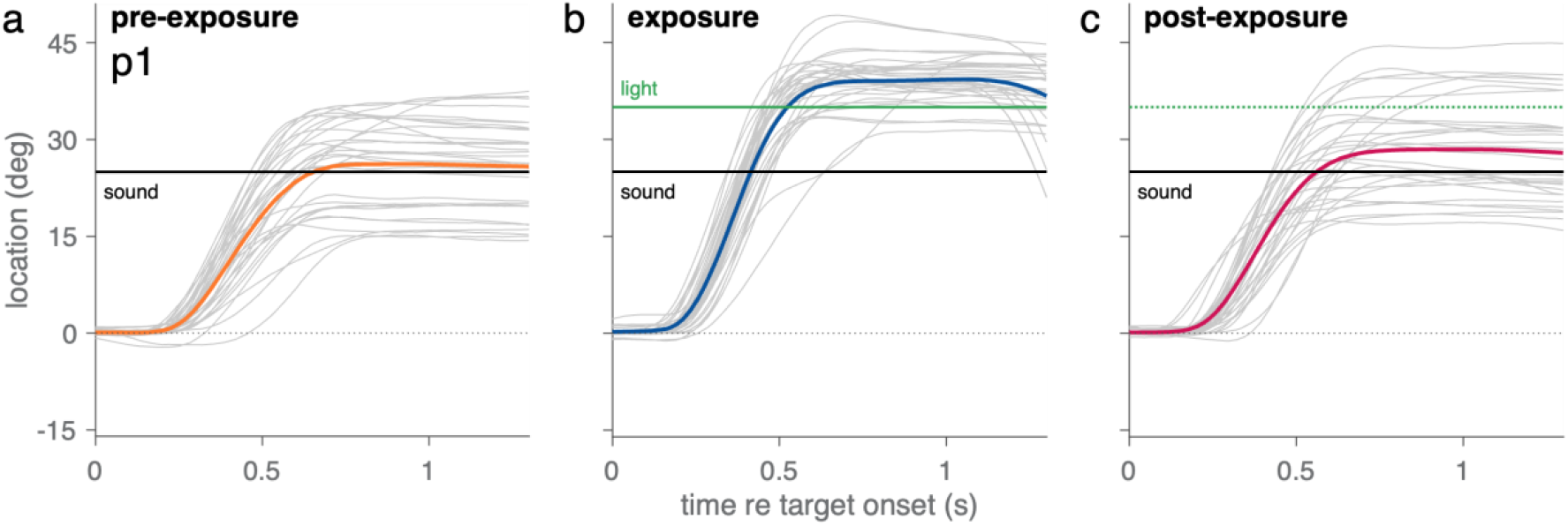
Head-orientation responses across experimental phases. Data shown are from a single participant (P1) during the three phases of one session (exposure sound: 5040 Hz). a) Pre-exposure phase. Head orientation traces (grey) in response to eight different test sounds presented at 25° (black line). Final head positions cluster around the sound location. The average response trace (orange) shows accurate localisation at the end of the rapid head orienting movement. b) Exposure phase. Head orientation traces to repeated presentations of a single narrowband sound (centre frequency: 5040 Hz) at 25°, paired with a visual distractor at 35° (green line). The average trace (blue) overshoots the sound location, ending near the light at 39°, indicating a ventriloquism effect. c) Post-exposure phase. Head orientation traces in response to the eight test sounds at 25°, now presented without the light. The average trace (pink) is biased toward the former visual distractor location, indicating a ventriloquism aftereffect.

## Data Analysis

### Modelling approach

Participants were instructed to localise band-limited noise bursts by orienting their head toward the perceived sound source. In the pre-exposure phase, sounds were presented in isolation, in complete darkness, and without any visual stimulus, allowing us to assess baseline sound-localisation behaviour. For each participant, session, and test frequency, sound-localisation responses were well described by a linear relationship between the true target location (T) and the response (R), characterised by a slope (gain) and a small intercept (bias).

We first examined whether pre-exposure localisation biases showed systematic structure across sessions or predicted post-exposure shifts in localisation. Sessions were conducted on separate days, with intervals ranging from one day to several weeks. We observed no systematic increase in pre-exposure bias across successive sessions (cf. ^26^). Moreover, biases in the pre-exposure phase did not predict localisation biases in the post-exposure phase (cf. ^27^). Baseline biases were generally small and unsystematic across conditions. Given this, and to facilitate stable estimation of localisation gain, all responses within a session were corrected for the average baseline response per participant, session, and test frequency. After this correction, auditory responses (*R*_*S*_) were well approximated by a linear function passing through the origin and proportional to the true target location (*T*; Fig. 3, blue). We therefore modelled sound-localisation responses as:

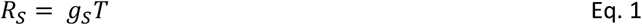

Here, *g*_*S*_ is the target-response or sound-localisation (S) gain (the slope of the target-response function). The gain is unitless and reflects how accurately perceived location tracks true location, with 1 reflecting perfect accuracy and 0 reflecting an absence of a relationship. With eight sound frequencies and twelve participants, this results in 96 gain estimates. Rather than estimating a separate gain for each frequency and participant, we expressed each gain *g*_*S*_ as a linear combination of fixed and random effects:

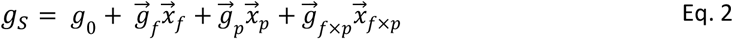

Here, *g*_*o*_ is the grand average gain, *g*_*f*_, *g*_*p*_, and *g*_*f×p*_ are the effects of frequency, participant, and their interaction, *x*_*f*_, *x*_*p*_, and *x*_*f×p*_ are the corresponding design vectors (e.g., indicator variables or contrast codes). These components reflect deviations from the grand average *g*_*o*_ due to frequency-specific and participant-specific effects, or their interaction. To illustrate the model, Fig. 3 shows data from example participant P2 for two sound frequencies: 793 Hz and 2000 Hz. The participant localised the 793-Hz sound with an overshoot (*g*_*S*_ = 1.34; Fig. 3a, yellow). This estimate decomposed as: an average gain *g*_*o*_ = 1.29, a frequency effect *g*_*f*_ = +0.07, a participant effect *g*_*p*_ = -0.12 and an interaction effect *g*_*f*×*p*_ = +0.1. For the 2000-Hz sound, the same participant showed accurate localisation with a slight undershoot (*g*_*S*_ = 0.93; Fig. 3b, yellow). This was decomposed into: *g*_*o*_ = 1.29 (same), *g*_*f*_ = -0.17, *g*_*p*_ = -0.12 (same), and *g*_*f*×*p*_ = -0.08. In both cases, the best-fit line (yellow) closely matched the observed data (yellow dots).

**Figure 3.**
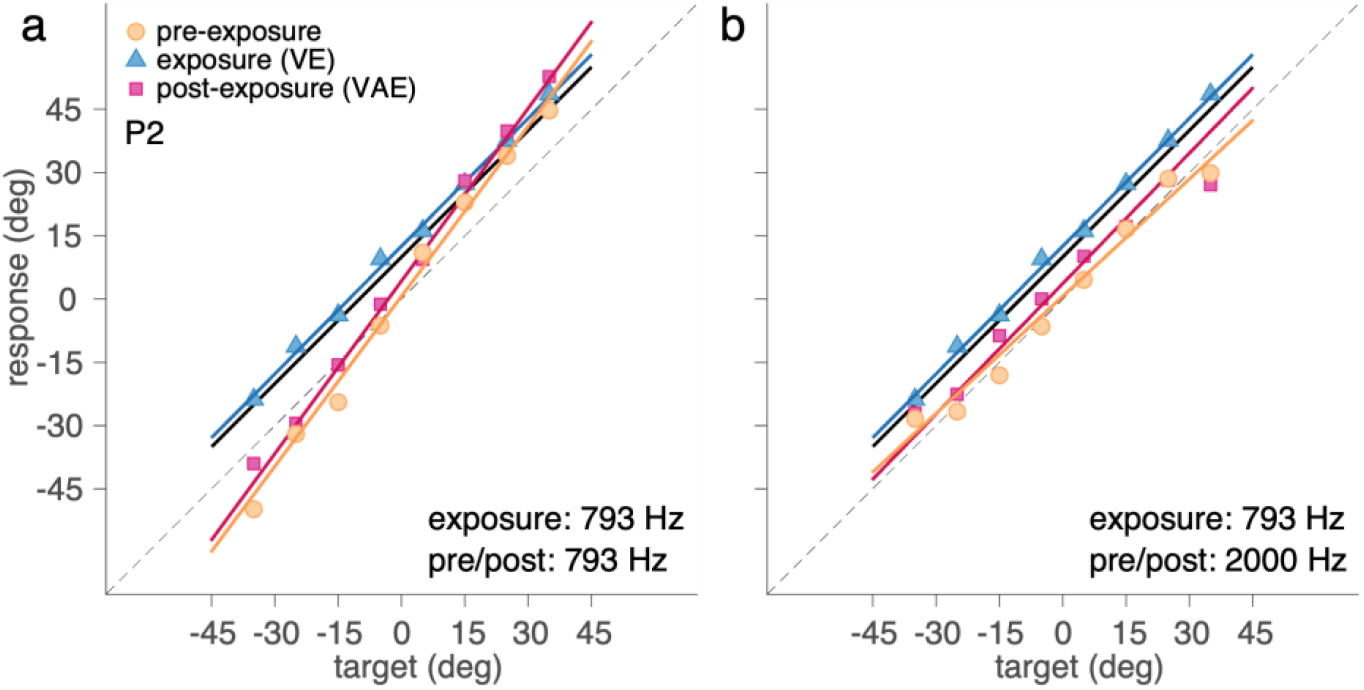
Example data and model fits. Target-response relationships for two sounds during the pre-exposure, exposure, and post-exposure phases. Panels show data from one participant (P2) for test sounds with a centre frequency at **(a)** 793 Hz and **(b)** 2000 Hz. In both cases, the exposure sound was a 793-Hz sound. Individual responses are shown as orange dots (pre-exposure), blue triangles (exposure), and pink squares (post-exposure); lines represent the corresponding model fits. The solid black bold diagonal line marks the location of the visual stimulus presented during the exposure phase (10° offset from the sound). The dashed black diagonal line indicates perfect localisation of the sound target.

During the exposure phase (Fig. 3, blue), sound localisation was systematically influenced by a visual stimulus (*V*) displayed 10 deg to the right of the auditory target (*T*). To quantify this ventriloquism effect, we modelled the response (*R*_*VE*_) as a linear function of both the auditory target location and the audiovisual disparity (*D*):

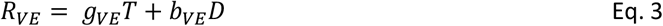

Here, *g*_*VE*_ is the ventriloquism-effect gain (slope) in the presence of a visual stimulus and *b*_*VE*_ is the ventriloquism-effect bias, a unitless parameter that reflects the proportion of the audiovisual disparity, *D*, that is incorporated into the participant’s response, with 1 reflecting a full bias towards the visual stimulus and 0 reflecting an absence of a visual bias. Importantly, this disparity *D* is not simply the physical 10° offset between the sound and visual stimulus, but rather the difference between the visual location *V* and the auditory response location:

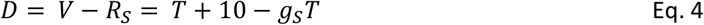

This definition of *D* centres the bias on the disparity between the visual stimulus and where the auditory target would have been localised in the absence of visual input (Eq. 1 and 2). With three exposure sounds and twelve participants, this results in 36 gain and 36 bias estimates. We expressed each gain and bias parameter as a linear combination of exposure sound (*e*), participant (*p*), and interaction effects, analogous to Eq. 2.

The slope and offset of the target-response line (as shown in Fig. 3, blue line) are the target-response gain and bias and can be derived from *g*_*VE*_, *b*_*VE*_, and *g*_*S*_ from Eqs. 3 and 4 as follows:

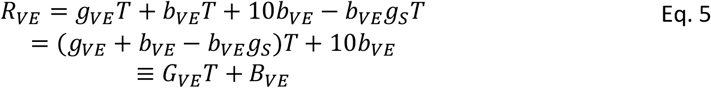

The target-response gain *G*_*VE*_ (dimensionless) and bias *B*_*VE*_ (in deg) during the exposure phase are then given by:

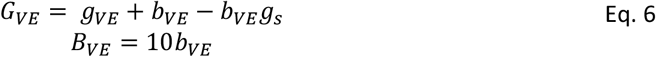

In this way, if *b*_*VE*_ is 0, *G*_*VE*_ reduces to *g*_*VE*_ and *B*_*VE*_ to 0 deg. If also *g*_*VE*_ = *g*_*s*_, sound localisation would not be affected by the visual stimulus, *R*_*VE*_ = *g*_*S*_*T* = *R*_*s*_. If *b*_*VE*_=1, *G*_*VE*_ reduces to *g*_*VE*_ - *g*_*s*_ + 1, and *B*_*VE*_ to 10 deg. If also *g*_*VE*_ = *g*_*s*_, sound localisation is completely dominated by the visual stimulus, *R*_*VE*_ = *T* + 10 = *V*. This analysis enables us to assess whether the presence of a simultaneous visual stimulus affects not only bias (as in the classical ventriloquism effect) but also the localisation slope, which may reflect changes in under- or overshooting behaviour. To illustrate, Fig. 3 (blue lines and triangles; note that panels a and b show the same exposure data) shows responses from participant P2 for the 793-Hz exposure sound. This participant was heavily influenced by the visual stimulus with a target-response gain near 1 (*G*_*VE*_ = 1.0) and a bias near 10 deg (*B*_*VE*_ = 12 deg). This indicates that the participant responded to the visual distractor, seemingly fully ignoring the auditory target.

To quantify the ventriloquism aftereffect, we modelled the response (*R*_*VAE*_) in the post-exposure phase as a linear function of the auditory target location and the remembered visual-auditory disparity (D) from the preceding exposure phase:

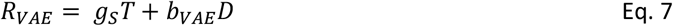

Here, *g*_*S*_ is the sound-localisation gain, shared across the pre- and post-exposure phase for all three sessions. The aftereffect bias *b*_*VAE*_ is a unitless proportion indicating the extent to which the response remains biased toward the location of the previously presented visual distractor, despite the absence of visual input during the post-exposure phase. A value of 0 indicates no aftereffect, while a value of 1 would indicate a complete shift equivalent to the prior visual offset. The gain and bias are estimated following the form of Eq. 2: *g*_*S*_ for each test frequency (*f*) and participant (*p*), and *b*_*VAE*_ for each test frequency (*f*), exposure sound (*e*), and participant (*p*). With eight test frequencies, three exposure sounds, and twelve participants, this results in 96 gain estimates (*g*_*S*_), and 288 bias estimates (*b*_*VAE*_).

Importantly, *g*_*S*_ was estimated using both pre- and post-exposure data, and we did not model a separate ventriloquism-aftereffect gain (as for the ventriloquism effect). This choice was motivated by the absence of any visual stimulus in both the pre-exposure (sound-localisation) and post-exposure (aftereffect) phases, and is consistent with empirical evidence showing that sound-localisation gain is not affected by the ventriloquism aftereffect^28^. In addition, inspection of the present data revealed no systematic differences in localisation gain between phases or across sessions, supporting this pooling strategy and allowing for more stable parameter estimates. To illustrate the model, Fig. 3 (pink lines and squares) shows post-exposure data from participant P2. For the 793-Hz test sound (Fig. 3a), the response was biased toward the location of the prior visual stimulus, with *B*_*VAE*_ = 4.3 deg. For the 2000-Hz sound (Fig. 3b), the aftereffect was lower (*B*_*VAE*_ = 2.2 deg) than that of the 793-Hz sound (Fig. 3a).

The target-response gain *G*_*VAE*_ (dimensionless) and bias *B*_*VAE*_ (in deg) during the exposure phase can be derived from Eq. 7 as follows:

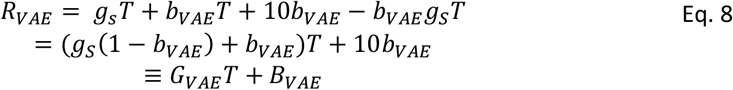

which yields:

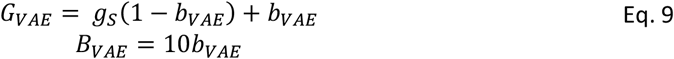

In this way, if *b*_*VAE*_ is 0, *g*_*VAE*_ reduces to *g*_*s*_ and *B*_*VAE*_ to 0 deg. This would imply there is no ventriloquism aftereffect, *R*_*VAE*_ = *g*_*s*_*T* = *R*_*S*_. If *b*_*VAE*_ = 1, *G*_*VAE*_ reduces to 1, and *B*_*VAE*_ to 10 deg. This would imply that the ventriloquism aftereffect, *R*_*VAE*_ = *T* + 10 = *V*.

### Bayesian Statistical Analysis

We used Bayesian inference to estimate model parameters from the observed data. Bayesian methods were chosen because they provide full posterior distributions for each parameter, offering both point estimates (e.g., means or medians) and credible intervals that quantify the uncertainty associated with these estimates. This approach allows for flexible implementation of complex hierarchical models, enabling simultaneous estimation at multiple levels.

The study employed a within-subject design with a large number of repeated trials per condition (hundreds per participant for the ventriloquism effect, dozens per participant for the ventriloquism aftereffect), and parameters were estimated using hierarchical models with partial pooling across participants and conditions. This structure increases statistical efficiency and stabilizes parameter estimates relative to analyses based on participant-level summary statistics alone.

All continuous variables were z-scored prior to analysis to facilitate parameter estimation. We applied standard normal priors (mean = 0, SD = 1) to the model parameters. These priors are weakly informative and appropriate for standardized data. To help the model converge, we based the prior mean for the ventriloquism bias on the mean effect observed in prior analyses. To account for individual variability and improve generalisability, all factors in the model (including test frequency, exposure frequency, and participant) were estimated using a hierarchical structure, allowing for partial pooling across levels of each factor. This approach reduces overfitting and provides more stable parameter estimates, particularly in conditions with sparse data. To aid interpretation, posterior estimates from the standardized model were back-transformed to the original scale using the means and standard deviations of the raw data, restoring units that are meaningful in the context of the observed behaviour.

Model fitting was performed via Markov Chain Monte Carlo sampling, in MATLAB (Mathworks, version R2021b), using the ‘matjags’ interface^29^ for JAGS^30^. Convergence was assessed using standard diagnostics, including the Gelman-Rubin statistic (all Rhat < 1.1)^31^ and effective sample size. To evaluate model fit, we performed posterior predictive checks, comparing model-generated data to observed patterns.

### Statistical inference

Statistical inference was based on Bayesian estimation of posterior distributions rather than null-hypothesis significance testing. Results are summarized using both point estimates (posterior means) and interval estimates in the form of 95% highest-density intervals (HDIs). Point estimates represent the most probable values of the parameters given the data, while HDIs quantify the associated uncertainty by indicating the range of parameter values with the highest posterior probability. The 95% HDI is defined as the narrowest interval containing 95% of the posterior probability mass, such that values outside the interval have lower posterior density than those within it.

Effects whose 95% HDIs did not include zero were interpreted as credibly different from zero. This Bayesian framework avoids dichotomous significance testing and instead emphasizes estimation of effect magnitude and precision. Differences between parameters were assessed using posterior mean differences and their corresponding HDIs, allowing direct probabilistic statements about the size and uncertainty of observed effects.

### Sample size and power considerations

Sample size was determined based on previous ventriloquism-aftereffect studies using similar within-subject psychophysical designs, which typically include 3-26 participants (e.g. ^5,10,12,14,32-35^). Given the large number of repeated trials per participant and the use of hierarchical Bayesian modeling, statistical power in the present study is driven primarily by the precision of within-participant estimates rather than by sample size alone. We did not perform an a priori power analysis in the classical frequentist sense, as the primary goal was estimation of effect sizes and generalisation patterns rather than binary hypothesis testing. Importantly, posterior uncertainty around the estimated ventriloquism effect and aftereffect parameters were small, as reflected by the narrow 95% highest-density intervals (e.g. Figs. 4 and 5). This indicates that the effects are estimated with high precision, driven by the large number of repeated trials per participant and the hierarchical modeling framework. In this context, inferential strength is reflected in posterior precision rather than in classical power considerations. Nevertheless, we acknowledge that the modest sample size limits sensitivity to subtle between-participant effects and that future studies with larger samples could further refine estimates of individual variability.

## Results

To examine how the ventriloquism aftereffect generalises across sound frequencies, participants localised sounds at various horizontal positions using head movements. Each participant completed three experimental sessions, each consisting of a pre-exposure, exposure, and post-exposure phase. During the exposure phase, one of three sounds—a low-frequency, high-frequency, or broadband sound—was paired with a visual stimulus located 10° to the right of the auditory target. Each of these sounds was tested in a separate session. In the pre- and post-exposure phases, participants localised eight sounds— seven narrowband and one broadband—without any visual stimulus. This design allowed us to assess baseline sound-localisation performance and determine the presence and generalisation of the ventriloquism aftereffect. Specifically, we tested whether the aftereffect was limited to the sound presented during exposure, generalised within cue types (low-frequency ITD vs. high-frequency ILD), or extended across the entire frequency spectrum.

We begin by presenting results on the ventriloquism effect - sound localisation in the presence of a simultaneous visual distractor - followed by the ventriloquism aftereffect, which reflects localisation shifts following prior exposure to audiovisual disparity. For both effects, we also report baseline sound-only localisation behaviour to characterize cue reliability.

### The ventriloquism effect

To assess the effect of a simultaneous visual stimulus on sound localisation, we analysed response gains and visual biases for the three sounds used during the exposure phases (793 Hz narrowband, 5040 Hz narrowband, and broadband). Figure 4a shows the target-response gains in the pre-exposure phase (*g*_*s*_; orange circles) and during the exposure phase (*G*_*VE*_; purple squares). In the pre-exposure phase (sound only), participants localised broadband sounds with high accuracy, yielding an average gain of 1.16 [1.14, 1.17]. In contrast, they exhibited overshooting for the narrowband sounds, with gains of around 1.3 [1.29, 1.31]. During the exposure phase (sound plus light), target-response gains were more accurate, being on average 1.03 [1.03, 1.04]. This high accuracy (gain close to 1) suggests that the presence of a visual stimulus improved localisation accuracy for narrowband sounds.

**Figure 4.**
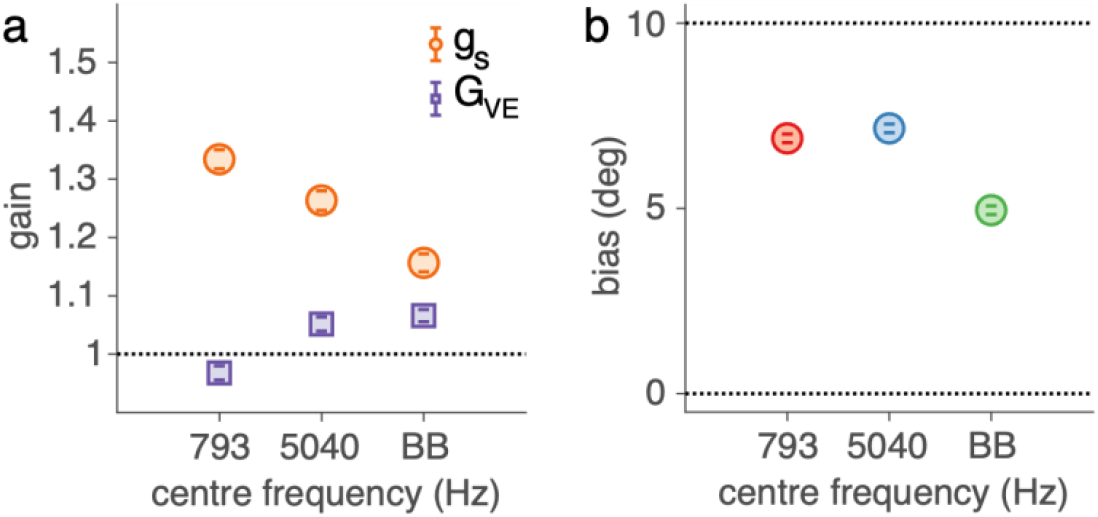
Ventriloquism effect. **a)** The sound-localisation response gains during the pre-exposure phase (g_s_, in orange) and the exposure phase (G_VE_, in purple). **b)** The bias (B_VE_) in deg for the three exposure sounds during the exposure phase. The mean and 95%-HDI of the posterior distribution on these parameter estimates are depicted as coloured circles and vertical error bars.

We also observed a robust bias (*B*_*VE*_) toward the visual distractor (Fig. 4b) in the exposure phase. The average bias across all sounds was 6.3 [6.3, 6.4] deg. This bias was stronger for narrowband sounds (7 [6.9, 7.1] deg) than for broadband sounds (5 [4.8, 5.1] deg). These results confirm a clear ventriloquism effect: a simultaneous light biased sound localisation, especially for narrowband stimuli.

### The ventriloquism aftereffect

Next, we analysed sound-localisation gains in the pre- and post-exposure phases to characterize how localisation responses varied with sound frequency before and after visual exposure. Specifically, we estimated the pre-exposure gain (*g*_*s*_; orange circles) and the post-exposure gain (*G*_*VAE*_; purple squares) for the eight test sounds (Fig. 5a). Both *g*_*S*_ and *G*_*VAE*_ showed a strong dependence on sound frequency. During the post-exposure phase, broadband sounds were localised with high accuracy, yielding a gain of 1.17 [95% HDI: 1.15, 1.18]. Narrowband sounds showed a characteristic V-shaped pattern, with the highest gains for the lowest and highest centre frequencies, peaking at 1.46 [1.42, 1.47] for the 500 Hz sound, consistent with overshooting, and the lowest gain for the 2000 Hz sound (1.12 [1.11, 1.14]). Importantly, the gains in the post-exposure phase (*G*_*VAE*_) slightly but systematically differed from the pre-exposure gains (*g*_*S*_). As explained in the Methods, this difference reflects the influence of a bias component (Eq. 8). In other words, the observed changes in gain during the pre- and post-exposure phase indirectly reflect the ventriloquism aftereffect.

**Figure 5.**
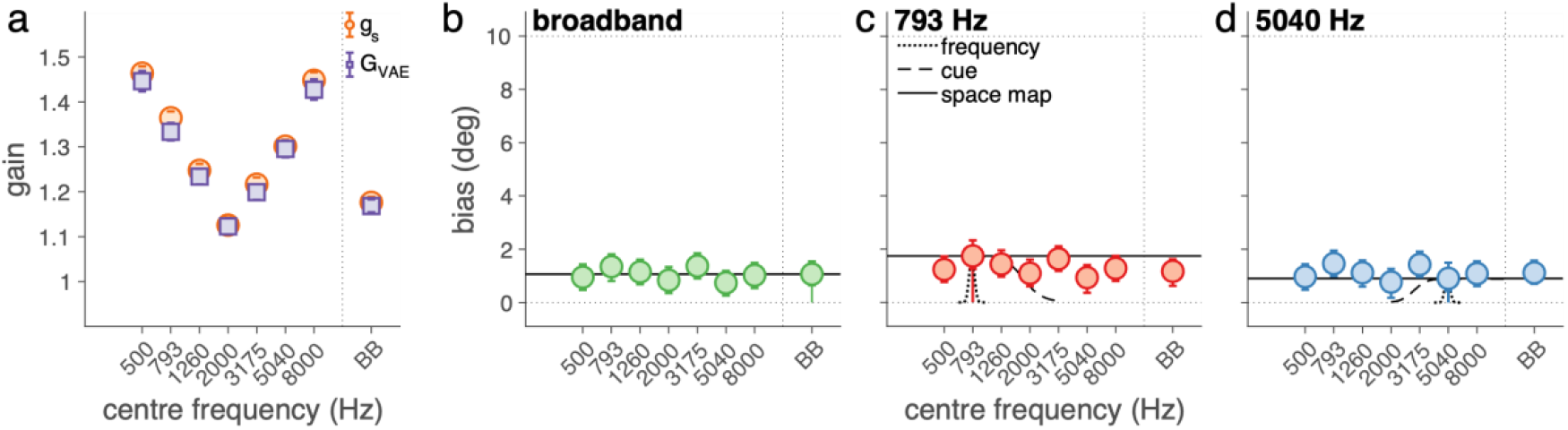
Ventriloquism aftereffect. **a)** The sound localisation gains, g_s_, pooled over pre- and post-exposure data for the eight test sounds, and the derived target-response gains, G_VAE_. **b-d)** The ventriloquism aftereffect bias, B_VAE_, for the **b)** broadband (BB), **c)** low-frequency (793 Hz), and **d)** high-frequency (5040 Hz) exposure sound as a function of test sound frequency. The mean and 95%-HDI of the posterior distribution on these parameter estimates are depicted as coloured circles and vertical error bars.

We analysed this aftereffect more directly by quantifying the bias term *B*_*VAE*_, which reflects the difference in perceived sound location between the pre- and post-exposure phases (Figs. 5b-d). The biases were relatively consistent, between 0.7 and 1.7 deg, suggesting that the aftereffect generalised beyond the specific exposure frequency to some extent. The magnitude of the aftereffect was similar across exposure sounds: 1.2 [1.0, 1.3] deg. Crucially, for the narrowband exposure sounds (Figs. 5b, c), a considerable aftereffect was present even for test sounds with a centre frequency far from the exposure centre frequency (e.g., the bias for the 8000-Hz test sound after 793 Hz exposure was 1.3 [0.8, 1.7] deg, Fig. 5c). This suggests that exposure to a low-frequency sound did not selectively bias low-frequency test sounds more than others, and similarly for high-frequency exposure. Taken together, these results show that the ventriloquism aftereffect, while small, is robust and broadly generalisable across the tested frequency range.

## Discussion

### Overview

This study investigated how the ventriloquism effect and its aftereffect depend on the spectral properties of sounds, to test whether these audiovisual interactions arise within frequency-specific auditory pathways or from a shared, frequency-independent spatial map. Participants localised sounds using head movements across three experimental phases: a baseline (pre-exposure), an audiovisual exposure (ventriloquism), and a post-exposure (aftereffect) phase. Each exposure phase involved one of three sounds, low-frequency, high-frequency, or broadband, paired with a consistent visual offset, while the aftereffect was probed using eight different sounds spanning a wide spectral range.

We found that sound localisation with narrowband noise was inaccurate and frequency-dependent, with overshoots particularly at low and high frequencies. Broadband sounds were localised accurately, presumably due to the presence of more reliable spatial cues. During the exposure phase, the ventriloquism effect varied across the three exposure sounds: broadband sounds showed the smallest bias, consistent with their higher spatial reliability. Moreover, the ventriloquism effect impacted not only bias but also the target-response gain, indicating that the presence of a visual stimulus alters both perceived direction and the slope of the auditory spatial response. The aftereffect was evident as a lasting bias in sound localisation, even in the absence of visual input. Importantly, the aftereffect generalised broadly across frequencies, regardless of whether the exposure sound was low-frequency, high-frequency, or broadband. That is, adaptation to an audiovisual discrepancy in one frequency band influenced localisation responses to sounds across the spectrum.

### Inaccuracy of narrowband sound localisation

A key feature of our study is the precise quantification of localisation accuracy for narrowband sounds. Consistent with earlier tone-localisation studies^36-40^, our data revealed large, frequency-dependent deviations between the perceived and actual locations of narrowband stimuli (Figs. 4a, 5a, orange). In contrast, broadband sounds were localised with high accuracy, showing only a slight overshoot. This discrepancy likely reflects differences in the reliability of spatial cues available for each sound type. Narrowband sounds primarily provide information through either interaural time differences (ITDs, e.g., 793 Hz) or interaural level differences (ILDs, e.g., 5040 Hz), but not both. While these cues support horizontal localisation, the resulting gains deviated substantially from unity by approximately 29%, highlighting a consistent overshoot. Additionally, narrowband sounds offer poor spectral content, rendering spectral elevation cues unreliable and making it impossible to resolve front-back or elevation ambiguities, often referred to as the ‘cone of confusion’^41^. In contrast, broadband sounds provide rich spatial information, including ITDs, ILDs, and spectral shape cues, enabling the auditory system to form a more precise and reliable spatial estimate. Thus, the lower localisation accuracy for narrowband sounds is best understood because of reduced cue reliability and ambiguity in spatial encoding.

A key feature of our study is the precise quantification of localisation accuracy for narrowband sounds. Consistent with earlier tone-localisation studies^36-40^, our data revealed large, frequency-dependent deviations between the perceived and actual locations of narrowband stimuli (Figs. 4a, 5a, orange). In contrast, broadband sounds were localised with high accuracy, showing only a slight overshoot. This discrepancy likely reflects differences in the reliability of spatial cues available for each sound type. Narrowband sounds primarily provide information through either interaural time differences (ITDs; e.g., 793 Hz) or interaural level differences (ILDs; e.g., 5040 Hz), but not both. While these cues support horizontal localisation, the resulting gains deviated substantially from unity, by approximately 29%, indicating a consistent overshoot. In addition, narrowband sounds offer limited spectral information, rendering spectral elevation cues unreliable and preventing resolution of front-back or elevation ambiguities (the ‘cone of confusion’^41^). As a consequence, spatial estimates based on narrowband sounds are both less reliable and more ambiguous. In line with this interpretation, recent work has shown that overshoots (i.e., gains > 1) are systematically associated with increased trial-to-trial variability, reflecting reduced localisation precision^27,42^. From this perspective, the pronounced overshoots observed for narrowband sounds may reflect a behavioural coping strategy in response to reduced cue reliability and imprecise spatial encoding. Notably, however, it remains unclear why reduced precision would give rise specifically to overshoots rather than undershoots. In other sensorimotor systems, such as visually guided saccades, increased uncertainty is typically associated with systematic undershooting, which has been argued to be an optimal strategy under noisy conditions^43^. The origin and functional significance of overshooting in sound localisation therefore remain open questions.

### Mechanisms in the ventriloquism effect

All three exposure sounds elicited a pronounced ventriloquism effect, traditionally defined as a spatial bias in auditory perception toward a simultaneously presented visual stimulus. This effect was strongest for the two narrowband sounds and slightly weaker for the broadband sound (Fig. 4b). This pattern aligns with Bayesian sensory integration, whereby auditory and visual cues are combined according to their relative uncertainties, with the more reliable cue receiving higher weight^2,44-46^. Because broadband sounds provide rich spatial cues and are localised more accurately (see previous section), they are treated as relatively reliable and are therefore less influenced by the visual stimulus. Narrowband sounds, in contrast, provide more ambiguous and less reliable spatial information, resulting in a stronger visual contribution and a larger visual bias.

Interestingly, for narrowband sounds the ventriloquism effect was not limited to a spatial shift (bias) but also involved a change in gain (Fig. 4a), bringing responses closer to veridical localisation. A natural interpretation is that the presence of a visual stimulus increases the reliability, and thus the precision, of the combined audiovisual spatial estimate. Consistent with the observation that overshoots covary with response imprecision^27,42^, the pronounced overshoots characteristic of auditory-only localisation for narrowband sounds are reduced during audiovisual stimulation. Accordingly, the observed gain changes are compatible with vision improving the precision of auditory spatial estimates, particularly when the auditory signal itself is unreliable. Because narrowband sounds exhibit substantial auditory-only overshoots, the effective audiovisual discrepancy can vary with target location, leading to concurrent changes in both response gain and bias during the exposure phase. This dual modulation highlights that the ventriloquism effect, especially for unreliable auditory signals, cannot be captured by a simple fixed spatial shift.

In this context, it is also worth considering the role of stimulus intensity. The present study employed a fixed sound level (55 dB SPL) and a constant visual luminance, chosen to ensure comfortable audibility and clear visibility across participants. At these levels, both auditory and visual stimuli were reliably perceived, making it unlikely that limitations in stimulus strength constrained localisation performance or multisensory correspondence. Holding sound level and visual brightness constant allowed us to focus on spectral differences. Nevertheless, if changes in localisation gain during audiovisual stimulation indeed reflect changes in spatial precision, future studies could further test this interpretation by systematically manipulating sound level and visual luminance to directly probe how stimulus strength shapes localisation precision, multisensory integration, and recalibration.

### Mechanisms in the ventriloquism aftereffect

We next examined the ventriloquism aftereffect across different combinations of exposure and test sounds. On average, participants exhibited a moderate aftereffect, with post-exposure sound-localisation responses biased by approximately 12% of the perceived audiovisual disparity (Figs. 5b-d). This bias was comparable across exposure sounds and extended across the full range of tested frequencies, indicating a broad generalisation of the aftereffect. Notably, this pattern contrasts with predictions from computational models by Magosso et al.^22,23^ and several earlier studies ^4-6,9^, which suggested that at moderate sound levels (∼55 dB), frequency-specific adaptation should dominate. In our study, however, we observed generalisation across the frequency spectrum, indicating that sound level alone may not fully determine the extent of generalisation. Instead, the present findings support the view that the expression of the ventriloquism aftereffect reflects an interaction between multiple experimental factors, including stimulus reliability, sound level, exposure structure, and task demands. From this perspective, frequency specificity and generalisation should not be regarded as mutually exclusive outcomes, but rather as different manifestations of recalibration that emerge under different experimental conditions; an issue that future studies should address by systematically varying these parameters.

An interesting aspect of the present results is that the magnitude of the ventriloquism aftereffect (Figs. 5b-d) did not mirror differences in the size of the ventriloquism effect across sound types (Fig. 4b). While the ventriloquism effect itself was larger for narrowband than for broadband sounds, consistent with differences in cue reliability, the aftereffect was comparable across exposure sounds. This apparent dissociation between immediate audiovisual bias and longer-term recalibration has been noted in previous studies^47,48^ and suggests that the processes underlying the ventriloquism effect and the ventriloquism aftereffect are at least partly distinct. One interpretation is that, whereas the ventriloquism effect reflects reliability-weighted cue integration and is therefore sensitive to cue uncertainty, the ventriloquism aftereffect serves a different functional role. In particular, recalibration may aim to reduce persistent spatial discrepancies and improve long-term accuracy rather than to optimise momentary precision, rendering it relatively independent of cue reliability. This view is consistent with arguments and empirical findings suggesting that multisensory recalibration can occur independently of cue reliability^49^, as well as with earlier reports showing weak or absent coupling between ventriloquism effect strength and aftereffect magnitude^48,49^. From this perspective, the broadly similar aftereffects observed across exposure sounds in the present study are compatible with a recalibration mechanism operating at a higher-level spatial representation that is less sensitive to stimulus-specific reliability.

## Conclusion

The present results demonstrate that broad frequency generalisation of the ventriloquism aftereffect can occur under conditions of a consistent audiovisual spatial offset, even at moderate sound levels. These findings support the hypothesis that the aftereffect reflects adaptation within a multisensory spatial representation, where auditory and visual signals are integrated into a shared topographic map. The reduced susceptibility of broadband sounds to the ventriloquism effect suggests a contribution of priors on sensory uncertainty. Together with earlier reports of frequency-specific recalibration, these findings support the view that the extent of generalisation is determined by both acoustic and non-acoustic factors (including experimental context). To conclude, the ventriloquism aftereffect reflects adaptive changes at a higher-level, modality-independent spatial map.

## Author contributions

A.J.v.O.: Funding acquisition, Resources, Supervision, and Writing - review & editing.

M.M.v.W.: Conceptualization, Data curation, Formal analysis, Funding acquisition, Methodology, Resources, Software, Supervision, Visualization, and Writing - review & editing.

N.C.H.: Formal analysis, Investigation, Project administration, Software, Validation, Visualization, and Writing - review & editing.

R.E.: Formal analysis, Funding acquisition, Investigation, Project administration, Visualization, Writing - original draft, and Writing - review & editing.

## Acknowledgements

We thank Günter Windau, Ruurd Lof, and Stijn Martens for their valuable technical assistance. We are also grateful to the volunteers for their participation in these experiments. Lastly, we thank Sem van Lankveld and Joep Ritzen for their assistance in the research project.

## Data availability

The data are openly available in the Data Sharing Collection of the Donders Repository at https://data.ru.nl/login/reviewer-2479852460/BJL7DGXGPO2XV3BHR7SPJNTHPW7PQDFKNXT36CA

[This is a link to give reviewers access to the data. After publication, this link will be replaced by: https://doi.org/10.34973/gbq9-yd62 ]

## Additional Information

Competing Interests: The authors declare no competing interests.

